# *AMHY* and sex determination in egg-laying mammals

**DOI:** 10.1101/2024.04.18.588952

**Authors:** Linda Shearwin-Whyatt, Jane Fenelon, Hongshi Yu, Andrew Major, Zhipeng Qu, Yang Zhou, Keith Shearwin, James Galbraith, Alexander Stuart, David Adelson, Guojie Zhang, Michael Pyne, Stephen Johnston, Craig Smith, Marilyn Renfree, Frank Grützner

**Author notes:** Joint first authors (Linda Shearwin-Whyatt, Jane Fenelon). joint senior author (Marilyn Renfree, Frank Grützner).

## Abstract

The sex chromosomes of egg-laying mammals (monotremes), which lack the sex determining gene *SRY*^1^, evolved independently to those of all therian mammals^2,3^. Here we characterise the candidate monotreme sex determining gene, the Y-localised anti-Müllerian hormone gene (*AMHY*)^4,3^ and trace its expression during the period of sexual differentiation. Monotreme *AMHX* and *AMHY* gametologues have significant sequence divergence at the promoter, gene and protein level, likely following an original allele inversion in the common monotreme ancestor but retain conserved features of TGF-β molecules. Expression of sexual differentiation genes in the echidna fetal gonad were significantly different from that of therian mammals. *AMHY* expression was seen exclusively in the male gonad during sexual differentiation, whereas *AMHX* was expressed in both sexes. Experimental ectopic expression of platypus AMHX or AMHY in the chicken embryo did not masculinise the female urogenital system, a possible result of mammalian specific changes to AMH proteins preventing function in the chicken. Our results provide fundamental insight into the first steps of monotreme sex chromosome evolution and sex determination with developmental expression data strongly supporting *AMHY* as the primary male sex determination gene.

## Introduction

Monotremes are the most basal, extant mammalian lineage, having diverged from therian mammals (marsupials and eutherians) around 188 MYA^3^. Monotremata, consisting of platypus and four echidna species, have a multiple sex chromosome system with five X chromosome pairs in females and five X and Y chromosome pairs in the male platypus and five X and four Y chromosomes in the echidna^2,5^. Monotreme sex chromosomes evolved independently from and share no homology with, therian sex chromosomes and lack the sex determination gene, *SRY*^1,6,7^.

The transition from autosome to sex chromosome is initiated by the acquisition of a sex determining gene on an autosome^8^. In therian mammals, changes to the *SOX3* gene and regulatory region gave rise to the sex determining gene on the Y, *SRY*^9,10^. During sex chromosome differentiation stepwise recombination suppression, surrounding the sex determining gene, produces a punctuated pattern of sequence divergence, termed evolutionary strata^4,3^ with the sex determining gene in the oldest stratum. A range of genes have become sex determining genes throughout evolution and most commonly arise from genes with conserved roles in sexual development^11^. The search for the monotreme sex determining gene has been hampered through the lack of Y chromosome sequence information and access to developmental stages. However, chromosome level genome assemblies of male platypus and echidna have confirmed that a male-specific copy of the anti Müllerian hormone (*AMH*) gene is the most likely sex determining gene of monotremes^12,4,3^. This gene, and its X chromosome gametologue, are in the oldest stratum (S0) of the platypus and echidna sex chromosomes and is the only known gene in this stratum with a conserved role in vertebrate sexual development.

AMH is a secreted hormone of the TGF-β superfamily, and it functions as a sex determining or differentiation gene in mammals, birds, reptiles, amphibians and fish^13,14,15,16^. In therian mammals, *AMH* is expressed after the testis-determining *SRY* and *SOX9* genes, and hence is a “downstream” component of the testis differentiation pathway. Sertoli cells secrete AMH, which causes regression of the Müllerian ducts that would otherwise develop into the female reproductive tract^17,14,18^. AMH binds to the AMH receptor type 2 (AMHR2) via the TGF-β domain to activate intra-cellular signalling via a type I receptor and the Smad proteins 1, 5, 8 and 4 leading to altered gene expression^16^. Mutations in *AMH* or in *AMHR2* cause persistent Müllerian duct syndrome in males and premature reduction of the follicle pool in females^19^. In species which lack Müllerian ducts, such as zebrafish, Amh promotes male development by regulating germ cell accumulation and inhibiting oocyte development or survival^20^.

*Amh* has emerged as a master sex determinant gene in a growing number of non-mammalian vertebrate species. The first identified was a male-specific copy of *amh* on the Y chromosome (*amhy*) in a fish, the Patagonian pejerrey, *Odontesthes hatcheri* where *amhy* knockdown caused male to female sex reversal^21^. Male-specific *amh* sex determining genes have since been identified in several species of teleost fish species^22,23,24,25,26,27,28^. *amhy* has arisen by independent, and lineage-specific, duplications rather than from a shared ancestral event^27^. In chicken, the *AMH* gene is located on chromosome 28, a region which has repeatedly and independently been recruited to sex chromosomes in vertebrate species^29^. These include the ZZ/ZW sex chromosome systems of several anguimorphs and geckos *Uroplatus* leaf-tail and *Coleonyx brevis* with *Amh* as the potential sex determining gene in these species^30,31,32^.

Identification of candidate sex determining genes in monotremes has been extremely challenging due to limited access to early developmental stages, lack of Y chromosome data and inability to perform genetic manipulation. However, the availability of new genome information and a significant improvement in captive breeding, have allowed us to access accurately timed and sexed echidna fetal and pouch young samples for the first time in over 150 years^33,34^. In our study, comprehensive genetic analysis and structural modelling has shown that monotreme *AMHX* and *AMHY* differentiation in promoter, intronic and coding sequence was an early step in monotreme sex chromosome evolution, but both have retained the conserved features of TGF-β proteins. We highlight differences in gene expression during monotreme and therian sexual development and show that expression of *AMHY* occurs exclusively in the male echidna bipotential gonad throughout the period of gonadal sexual differentiation, strongly supporting the conclusion that *AMHY* is the primary sex determining gene of monotremes. Over-expression of AMHX or AMHY in an avian model system (chicken embryo) suggest that AMH receptor binding and signalling has undergone specific changes in mammals, explaining the lack of a gonadal phenotype seen in the developing chicken embryo.

## Results

### An inversion involving AMH in the monotreme common ancestor

We compared the genomic region around *AMHX* and *AMHY* for signs of differentiation. In monotremes the *AMH* gene is located on the sex chromosomes, in contrast to therian mammals in which it is autosomal. *AMHX* is located in the X specific region of X1 in platypus and echidna in the oldest stratum (Fig. 1A). Synteny (collinearity) of genes flanking *AMH* is conserved throughout vertebrate species including human chromosome 19 and chicken chromosome 28 with a chromosomal breakpoint flanking the *LING03* gene (Fig. 1A, blue arrowhead). The *SF3A2* gene lies immediately upstream the *AMH* gene (within 0.6 kb of the start site) with the *JSPR1* gene downstream. An Follicle-Stimulating Hormone-responsive enhancer involved in upregulation of *AMH* expression in Sertoli cells of the prepubertal testis lies 1.9kb upstream of human AMH gene overlapping with *SF3A2*^35^. This and conservation of the local chromosomal context of *AMH* across divergent vertebrate species suggests that sequences adjacent to AMH genes can influence AMH expression. *AMHY* is located in the non-recombining region of chromosome Y5 in platypus and Y3 in echidna. The local gene order surrounding the monotreme *AMHY* genes is different from the X copy, likely due to inversions and Y chromosome differentiation and may affect expression of *AMHY* compared to *AMHX*. The *AMHY* genes are flanked by the *STK11* and *PIAS4* genes in both platypus and echidna. These genes are both on chromosome 28 of chicken and 19 of human but are not adjacent, being 1.5MB and 2.7 MB apart respectively. In echidna, the *STK11* gene is 22MB from *AMHX* and *PIAS4* is on chromosome X2. Furthermore, the relative orientation of the *STK11* and *AMHY* genes is altered compared to *AMHX* and *STK11* on monotreme X1 and on chicken chromosome 28, suggesting inversion events involving the proto-*AMHY* may have occurred in the monotreme common ancestor (Fig. 1B).

**Figure 1.**
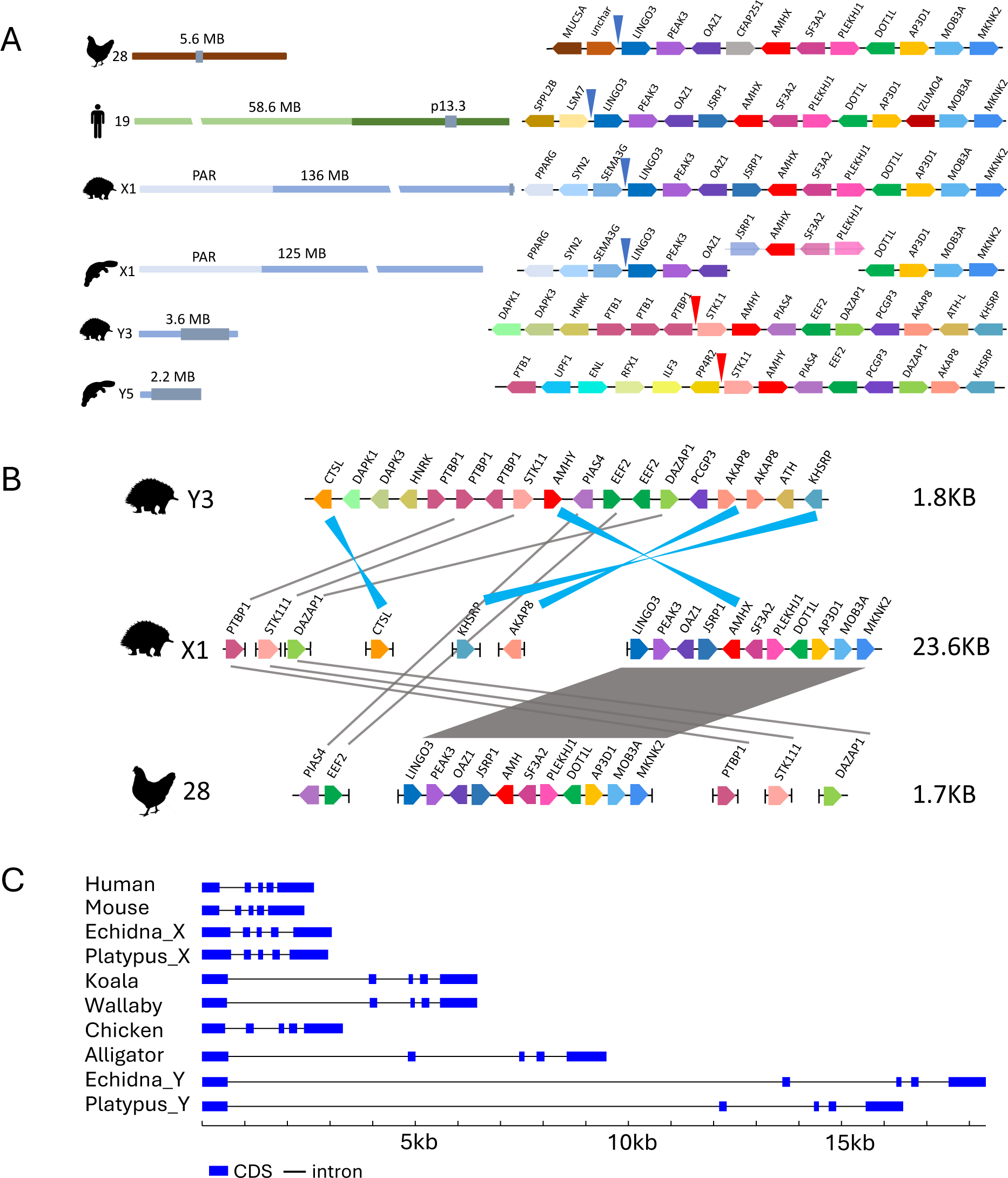
Synteny and gene structure analysis of monotreme *AMHX* and *AMHY*. (A) Synteny between chicken human and monotreme AMH genes. Chicken chromosome 28, brown; human chromosome 19, green with q arm in light green; monotreme chromosomes blue, with PAR in light blue; grey box syntenic region in expanded view showing genes as coloured directional boxes. Blue arrow, break in synteny with monotreme X chromosomes; red arrow, break in synteny between monotreme Y chromosomes. (B) Comparison of echidna genes flanking AMHX/Y and chicken chromosome 28. Genes as coloured directional boxes. Connecting lines grey, genes in same orientation, blue, genes in opposite orientation; Not to scale. (C) Gene organisation of monotreme *AMHX* and *AMHY* compared to that of the human, mouse, koala, wallaby, chicken and alligator *AMH*. Each gene contains 5 translated exons. CDS, coding sequence.

The monotreme *AMH*X and *AMHY* genes consist of 5 exons with similar organisation to that of other vertebrate *AMH* genes (Fig. 1C). The introns of the *AMHY* genes are considerably larger than those of the *AMHX* or *AMH* genes (Fig. 1C) largely as a result of repeat invasion as is often seen with Y chromosome gametologues (Supplemental Data Fig 1A) ^36,37^. Furthermore, repeat content is higher in the *AMHY* flanking regions compared to *AMHX*. 7kb upstream of monotreme *AMHX*, repeat coverage is just under 40% whereas the same region upstream of *AMHY* has 60 to 72% repeat coverage (data not shown). The *AMHY* genes also show specific examples of repeat invasion. For example, intron 1 has undergone large expansion due to LINE L2 and SINE intronic insertions in both platypus and echidna (Supplemental Data Fig 1A). One conserved LINE L2 insertion is present in intron 4 of the monotreme *AMHX* genes. Conservation of this L2 insertion in both platypus and echidna shows that this occurred before divergence of platypus and echidna and its conservation for over 55 million years suggests functional importance.

### *AMHX* and *AMHY* have diverged through sex chromosome differentiation

Comparative analysis of the monotreme *AMHX* and *AMHY* genes shows that the coding sequence of the gametologues have substantially diverged from each other sharing only ∼60% DNA identity (Supplementary Table 1). In contrast, there is greater between species similarity between the platypus and echidna *AMHX* genes and between the platypus and echidna *AMHY* genes (93% and 90% sequence identity respectively). This is also reflected in the nucleotide substitution rate (Supplementary Table 2). The Ka/Ks ratios of monotreme *AMHX* and *AMHY* (<1) compared to *AMH* of other vertebrates indicates that the genes are under purifying selection, implying that their functional/molecular attributes have been conserved.

At the protein level, the AMHX sequences are over 90% identical and the monotreme AMHY proteins are also highly similar with just under 90% identity. Importantly, the AMHX and AMHY proteins within each species, are just over fifty percent identical. This suggests that the divergence of the *AMHX* and *AMHY* gametologues which began in early monotreme evolution at around 100 MYA largely occurred prior to divergence of the echidna and platypus and has been maintained since then (Supplementary Table 3 and Fig. 2A). The monotreme AMHX and AMHY proteins have the conserved features of AMH proteins, including a predicted signal peptide, conserved cleavage site between the N-terminal and C-terminal domains and the 7 conserved cysteine residues of the TGF-β domain (Fig. 2A, Supplemental Data Fig. 1B). The TGF-β domain is the most highly conserved region of AMH proteins as it binds directly to the AMH receptor leading to signalling and this is also true of the monotreme AMHX and AMHY proteins (Supplementary Table 4, Fig. 2A). Within the monotreme TGF-β domains there are only two amino acid differences, but the protease cleavage sites differ between AMHX (RAQR) and to AMHY (RVQR) which may impact on cleavage into the biologically active domains (Fig. 2A).

**Figure 2.**
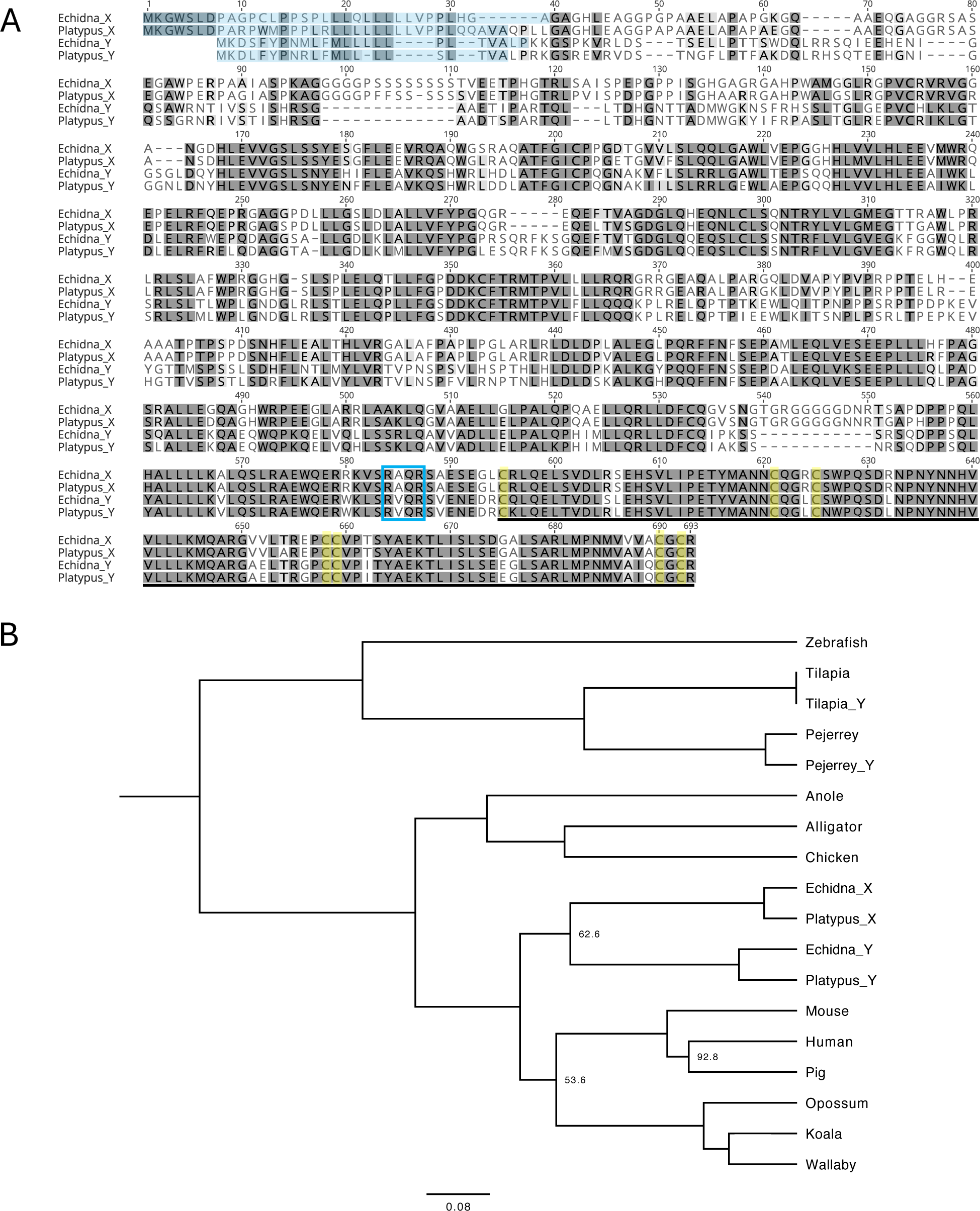
Comparative analysis of monotreme AMH proteins. (A) Alignment of the monotreme AMHX and AMHY proteins. Grey shading, amino acid similarity with identical residues (dark grey); blue shading, signal peptides; blue box, protease cleavage site; underline, TGF-β domain with conserved cysteine residues highlighted in yellow (B) Phylogenetic tree of vertebrate AMH proteins. Scale bar, branch length; branch labels, Bootstrap values shown if below 95%.

The structure of the TGF-β domains of AMHX and AMHY from both platypus and echidna were assessed by *in silico* modelling^38,39^. The best models obtained were all based on human AMH TGF-β domains with high confidence. As expected, based on the high degree of amino acid conservation, the predicted structures are very similar to each other with root mean square deviations of less than 0.2 Å across the full length (106 amino acids) of the TGF-β domain. The most variable regions were found in the loops at the distal tip of the structure, which interact with the AMHR2 receptor^40^. All known TGF-β family ligands form dimers which can interact with two subunits of the extracellular domain of the cognate receptor. Prediction of platypus AMHR2-AMHX and AMHR2-AMHY multimeric complexes of the same stoichiometry was done with AlphaFold multimer^39^, giving structures with topology very similar to the experimentally determined human AMHR2-AMH complex^40^ (Supplemental Data Fig. 1C). The predicted platypus AMHR2 extracellular domain showed conservation of the “3 finger toxin” fold characteristic of the extracellular ligand binding domain of the TGF-β family. The majority of the key contact residues essential for AMH signalling in the human AMHR2 receptor^40^ are conserved in the platypus receptor complexes (Supplemental Data Fig. 1D), suggesting that the monotreme AMHX and AMHY proteins both signal through the AMHR2 receptor.

### *AMHY* has novel transcriptional control elements

As evolution of new sex determining genes can be caused by alterations to regulatory regions which change the pattern of expression, we compared the proximal promoters of *AMHX* and *AMHY* to each other and to those of *AMH* genes of other mammalian species. Comparison of the human, mouse, tammar wallaby *AMH* and the monotreme *AMHX* and *AMHY* showed that the *AMHX* promoters are more similar to each other and to those of therian *AMH* genes than they are to the monotreme *AMHY* promoters (Fig. 3A and Supplemental Data Fig. 2).

**Figure 3.**
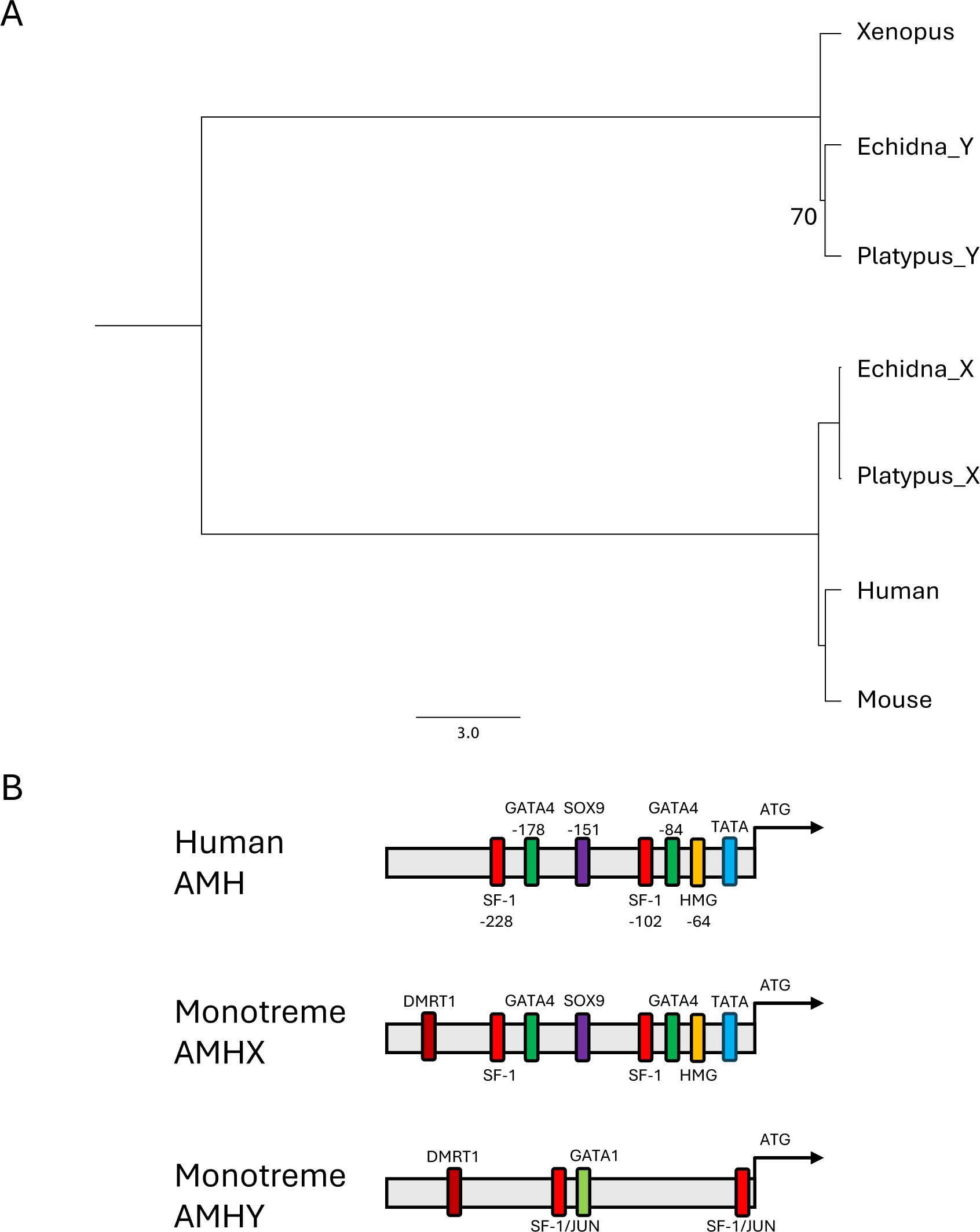
Comparative analysis of *AMH* proximal promoters. (A) Phylogenetic tree of vertebrate AMH promoters. Scale bar, branch length; branch labels, Bootstrap values shown if below 95%. (B) Schematic representation of the human *AMH,* monotreme *AMHX and AMHY* promoters showing transcription factor binding sites (not to scale). Binding sites for GATA1, GATA1 transcription factor;GATA4, GATA4 transcription factor; SF-1, steroidogenic factor-1; JUN, Jun; DMRT1, doublesex and mab-3 related transcription factor 1 ; SOX9, *Sry*-type box 9; HMG, High mobility group protein; TATA, TATA binding factor; arrow, +1 translation start site.

In humans, expression of AMH in Sertoli cells of the fetal testis is activated by direct binding of SOX9 to the proximal *AMH* promoter^41^, following its activation by SRY. Binding of steroidogenic factor-1 (SF-1), GATA4 binding protein (GATA4) and Wilms’ Tumour protein (WT1) to the proximal promoter enhance transcription of *AMH* in the fetal gonad (Fig. 3B). Putative binding sites for SOX9, GATA4, SF-1 and HMG in the monotreme *AMHX* promoters are conserved in sequence and relative position with those of the mammalian *AMH* promoters. In contrast, binding sites for SOX9, GATA4, SF-1 and HMG are not found in monotreme *AMHY* promoters. Rather, putative binding sites for SF-1/Jun, GATA1 and DMRT1 are conserved in the promoter region in both platypus and echidna *AMHY*. The putative DMRT1 binding site, corresponding to the DMRT1 binding site identified by the SELEX method^42^, is also present in the monotreme *AMHX* promoters (Fig. 3B and Supplemental Data Fig. 2). This shows that the *AMHY* promotor region has differentiated, and the changes have been conserved between *AMHY* in platypus and echidna suggesting that they are functional.

### *AMHY* is expressed in the male echidna gonad throughout sexual development

*In situ* hybridization was used to assess the cellular sites of expression of monotreme *AMHX* and *AMHY*, in adult gonads (Fig.4A, B; expression purple stain). *AMHX* was expressed in the granulosa cells of the follicles in both echidna and platypus (Fig. 4A) as is the case for therian AMH^43^. Both *AMHX* and *AMHY* were expressed in DMRT1/SOX9^+^ Sertoli cells of both platypus and echidna (Fig. 4B) as in therian mammals^44^.

**Figure 4.**
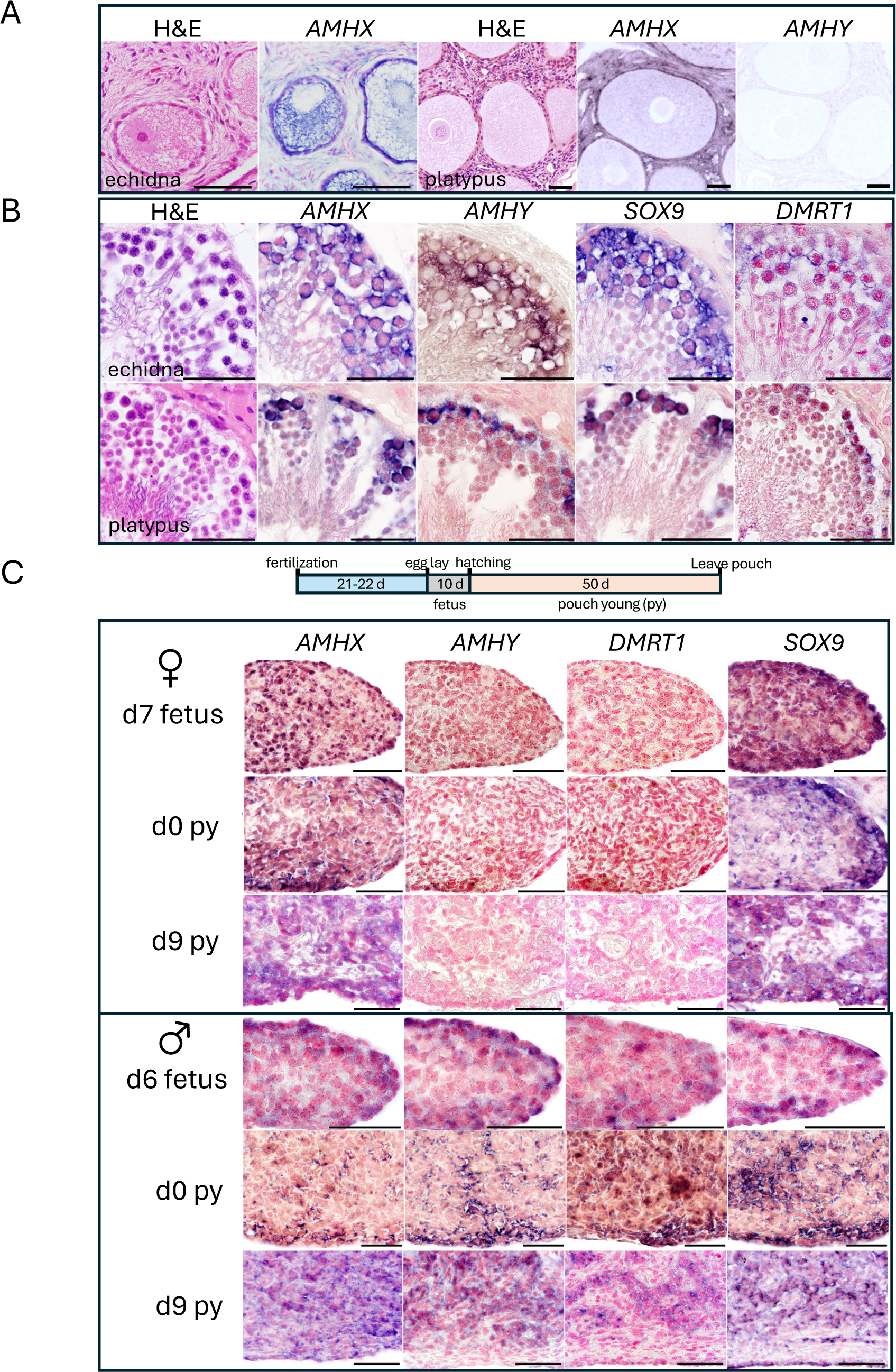
Expression analysis in monotreme gonads by *in situ* hybridization. Haematoxylin and eosin stained adult (A) ovary sections showing *AMHX* expression in the echidna and platypus and *AMHY* in the platypus. (B) testis sections showing *AMHX, AMHY, SOX9* and *DMRT1* expression in echidna and platypus testis. (C) Timeline of early echidna developmental stages (upper). Expression of *AMHX*, *AMHY*, *DMRT1* and *Sox9* in echidna female (middle) and male (lower) gonads during gonadal sexual differentiation. Scale bars: 50μm.

For a gene to be male sex determining it must be expressed in the male fetal gonad during the period of gonad differentiation. In echidna, the ovary and testis first become morphologically differentiated after hatching between the day 2 to day 4 of the pouch young (py) stage (JC Fenelon and MB Renfree unpublished results, Figure 4C). We assessed gene expression in developing echidna gonads from a day 6 fetus (in egg, where the day of egg laying is designated d0) through to day 11 post-hatching pouch young stage. The sex of all echidna fetuses and pouch young was determined genetically using published markers^45,31^ (Supplemental Data Fig. 3A and B). *In situ* hybridization shows *AMHX*, *AMHY, SOX9* and *DMRT1 expression* in the echidna fetal gonad throughout the period of sexual development (Fig. 4C) and specifically in the bipotential male gonad from the d6 fetus through to d0 py. By the d9 py, *AMHY* was clearly expressed in the Sertoli cells of the testis cords, similar to *DMRT1* and *SOX9*. *AMHX* and *SOX9* were also expressed in the developing female gonad at all stages examined but expression of *AMHY* and *DMRT1* were not detected (Fig. 4C). These results were confirmed by RT-PCR (Supplemental Data Fig. 3C). RT-PCR showed expression of *DMRT1* in female d2 samples but not in d4 samples. Genes important for therian AMH expression (*SF-1*, *WT1* and *GATA*) or function (*AMHR2*) were expressed from d0-d4 py in both the male and female gonads whilst *GATA4* was expressed in all except the male d0 py gonad (Fig. 5A). In contrast, *GATA1* was only present in the d2 and d3 py male gonads (Fig. 5A). Expression of the *AMHR2* in the bipotential gonads of both sexes suggests that *AMHX* and *AMHY* signalling is required throughout this period of development. Importantly, expression of *AMHY* in the male, but not female, gonad throughout the period of gonadal specification supports its role as the primary male sex determining gene. These data also indicate that *GATA1* and *DMRT1* maybe involved in monotreme male sex determination (summarized in Fig 5B).

**Figure 5.**
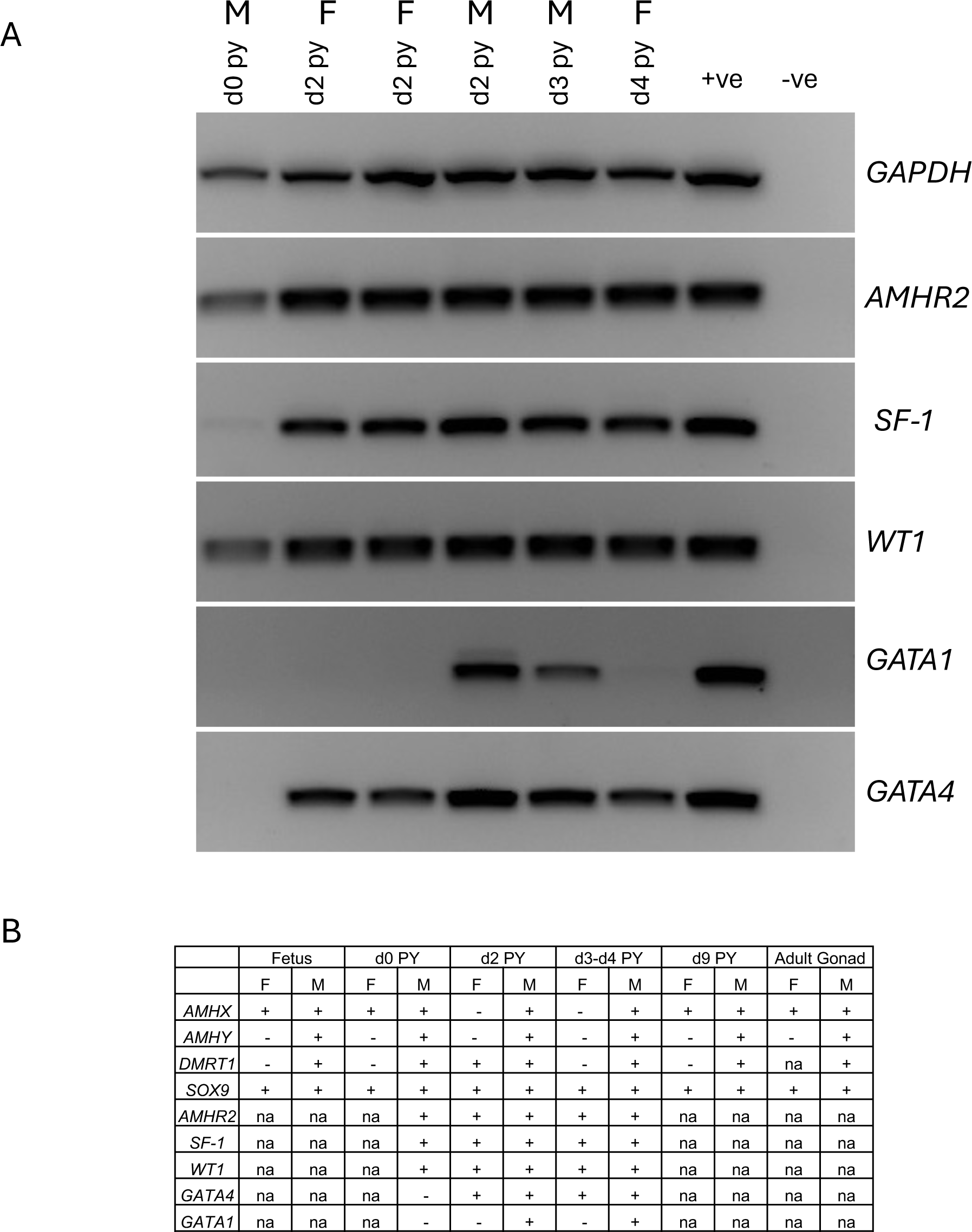
RT-PCR Echidna pouch young gene expression analysis. (A) Expression of *AMHR2*, *SF-1*, *WT1*, *GATA1* and *GATA4* with *GAPDH* as loading control in pouch young gonads from d0-d4 py with either adult testis or ovary as a positive control. *GAPDH*, *AMHR2, SF-1 and GATA4* were expressed in both sexes at all stages examined whereas GATA1 was only expressed in the d2 and d3 py males. py: pouch young (B) Summary table of echidna gene expression. +, expression; -, no expression; PY, pouch young; F, female; M, male; py, pouch young; na, data not available.

### Ectopic expression of *AMH* proteins during sexual development in a non-mammalian model system

We investigated whether AMHX and AMHY proteins can affect sexual development in a model animal, the chicken. In chicken embryos, forced over-expression of AMH in female embryos can masculinise the gonads, while knockdown of the gene blocks proper gonad formation in both sexes^46,47^. AMH and AMHR2 are expressed in both sexes of the chicken from early embryonic stages, prior to sexual differentiation^48,49^. This early timing, together with the known masculinising effects of exogenous chicken AMH on avian gonads, makes the chicken embryo the most suitable model for assessing monotreme AMH function. Chicken embryos were infected at the blastoderm stages with viral vectors over-expressing either chicken AMH or platypus AMHX or AMHY (each GFP-tagged). Embryonic chicken urogenital systems were examined at E7.5 (stage 32). Over-expression of chicken AMH in the chicken embryo produced the expected phenotype, characterised by small elongated (male-like) gonads and regression of Müllerian ducts in both sexes (Supplemental Data Fig. 4) proving successful over-expression as previously described^47^. Over-expression of platypus AMHX or AMHY proteins did not produce a discernible phenotype in the developing gonads of either males or females (Supplemental Data Fig. 4) even with the chicken specific cleavage site required for processing into the functional AMH domains (panels labelled +CC). Detailed sequence analysis shows that chicken and monotreme AMH proteins differ in key residues at the interaction face required for human receptor binding, as defined recently for the human AMH/receptor interaction^40^ (Supplemental Data Fig. 5A). Comparison of the TGF-β domains of AMH across taxa indicated that the residues implicated in binding and/or activity of human AMH and AMHR2 are completely conserved in the eutherian mammal species examined and largely conserved within all three mammalian groups (Supplemental Data Fig. 5B). This suggests that changes in the AMH/AMHR2 system in early mammalian evolution may prevent mammalian AMH proteins functioning through the chicken receptor.

## Discussion

The divergence of monotremes from therians predates the evolution of *SRY* as the sex determination gene in therian mammals. Monotremes evolved a complex XY sex chromosome system that shares no homology with therian sex chromosomes. Instead, these chromosomes have homology to the avian Z sex chromosome which includes the conserved sex determining gene, *DMRT1* on an X chromosome^50,2,51^. The absence of a *DMRT1* gametologue on the Y chromosomes and its localisation in stratum S3 of the X chromosomes, excludes *DMRT1* as the primary monotreme sex determination gene^3^.

Identification of Y-specific *AMH* gametologues in the oldest stratum of the monotreme sex chromosomes (platypus Y5, echidna Y3^4,3^) makes *AMH* the frontrunner as the monotreme sex determination gene. Recent estimates suggest that monotreme sex chromosomes evolved at least 100 MYA and that the differentiation of the *AMH* gametologues occurred early in the evolution of the monotreme sex chromosome complex (Y Zhou *et al*., submitted 2024). This is fundamentally different to examples in fish where *Amh* has become a sex determining gene by duplication and translocation onto the future or existing Y chromosome^52^ and can maintain a high level of sequence identity with autosomal *Amh*^23^. By contrast monotreme *AMHX*/*AMHY* are highly differentiated with the genomic context conserved for AMHX but not *AMHY*. Our data are consistent with an inversion of one *AMH* allele producing the proto-Y chromosome in the monotreme ancestor. Recombination suppression caused by the inversion, would be a first step in sex chromosome differentiation. Conserved changes in transcription factor binding sites in the proximal regulatory region of *AMHY* and loss of the upstream *SF3A2* gene, which impacts *AMH* expression in therian mammals, further supports a male specific function. Accumulation of intronic transposable element insertions in *AMHY* may also affect expression of the gene compared to *AMHX*. This can be particularly important for developmental control genes where precise temporal control is required and large introns can cause transcriptional delay^53^. Interestingly most of the LINE L2 and SINE MON1 insertions are conserved between platypus and echidna suggesting that repeat invasion occurred in the common ancestor more than 55 MYA.

Phylogenetic analysis shows separation of the AMHX and AMHY proteins, signifying evolutionarily conserved differences in the gametologues. Both proteins retain the features of functional TGF-β family members with subtle substitutions that may affect signalling and proteolytic cleavage. Together these results provide evidence that the monotreme X- and Y-localised *AMH* genes encode functional signalling molecules with significant differences in both coding and regulatory sequence which may have conferred a sex determining function on *AMHY*.

Similar to other mammals, *AMHX* is expressed in the granulosa cells of the ovary and Sertoli cells of the testis while *AMHY* expression is restricted to Sertoli cells of the testis. Importantly, in the developing echidna fetus *AMHX* and *AMHR2* are expressed at least from the day 6 fetal stage in male and female, about 8 days before the gonads are morphologically different. This is notably earlier than the onset of *AMH* expression in therian mammals. In the developing echidna male gonad, *AMHY* shows expression in all stages examined, from the earliest day 6 fetal stage to the day 9 pouch young stages, consistent with a role in monotreme gonad specification. In therians, *AMH* expression is upregulated in response to increased *SOX9* expression after gonad specification in the developing mouse testis by E12.5 when it results in loss of the Müllerian ducts in males^43^. Echidna *AMHY* expression is more similar to non-mammalian vertebrates (chicken, turtle, American alligator and bearded dragon, *P. vitticeps*) in which *Amh* expression also precedes that of *Sox9*^54,55,56,57^. Such co-expression of autosomal Amh and sex determining Amhy in the gonad during sexual differentiation is also seen in the Northern Pike (*Esox lucius*)^27^. We suggest that co-expression of AMHX and AMHY in the male gonad may be required to trigger male development, whereas female development would occur in the absence of AMHY. AMHX/Y heterodimers may result in altered signalling through AMHR2. In other examples where *amhy* is sex determining (eg *O. hatcheri*) knockdown experiments suggest that *amhy* represses the female pathway genes *foxl2* and *cyp19a1a* required for ovary development. *AMHY* could play a similar role in the monotremes, repressing expression of genes required for ovarian development.

In therian mammals, *SOX9* expression is low in the bipotential gonad of both sexes but is the first gene upregulated in the male gonad by SRY which causes upregulation of *AMH* expression^41^. It was therefore unexpected to see *SOX9* expression in the echidna throughout the sexual development period in both the female and male gonad. In therians SOX9, SF-1, GATA4 and WT1 directly activate *AMH* expression later during male gonad development^58^. Here we show that *SOX9*, *SF-1*, *GATA4* and *WT1* are expressed during the echidna gonadal differentiation period in both sexes. We also show that *DMRT1* has male biased expression in the bipotential echidna gonad from the earliest stage examined up until at least the day 9 pouch young, the latest stage examined, and *GATA1* is exclusively expressed in the male gonad together with *AMHY*. Conserved potential binding sites for GATA1 in the monotreme *AMHY* promoters and for DMRT1 in the promoters of monotreme *AMHX* and *AMHY* suggest that these factors may be involved in regulating *AMHY* expression during gonad specification in monotremes. These data highlight significant differences between mammalian sexual development pathways with *SRY* (therian) or without *SRY* (monotremes).

In chicken AMH and AMHR2 are expressed prior to gonadal differentiation and overexpression of chicken AMH causes masculinization of female gonads^47^. Over-expression of monotreme AMHX and AMHY in the chicken did not recapitulate the chicken AMH over-expression phenotype. We suggest that inefficient binding and/or signalling through the chicken AMHR2 may prevent activity of the monotreme AMHX and AMHY proteins. Interestingly chicken AMH can induce regression of mammalian (rat) Müllerian ducts, while rat AMH has no effect in chicken^59^. Analysis AMH proteins of vertebrate species suggests that the residues involved in AMHR2 binding by human TGF-β domain of AMH^40^ are only conserved in mammals suggesting that mammalian AMH proteins more broadly may not function in bird models.

Monotreme sex determination has been a mystery since the first unsuccessful attempts to identify the mammalian sex determination gene, *SRY* over thirty years ago. Here we provide a comprehensive analysis of the monotreme *AMHX* and *AMHY* genes and the first insight into the sexual development pathway of this mammalian lineage which lacks *SRY*. We show that the *AMHX* and *AMHY* gametologues are highly differentiated in genomic context and sequence likely caused by early events in the evolution of monotreme sex chromosomes. Importantly in the developing echidna gonad, *AMHX* is expressed earlier than *AMH* in therian mammals and *AMHY* is exclusively expressed in the developing male fetal gonad throughout the period of gonad specification, supporting a role for *AMHY* as the primary monotreme male sex determination gene. As such,, it would be the first example of a soluble growth factor acting as a sex determination gene in mammals.

## Methods

### Gene and Protein analysis

Platypus AMHX (NC_041749.1). Platypus AMHY Y5 (NC_053179.1): 1478061-1508837, geneID 114808765. Echidna AMHX X1(NC_052101.1):135469955-135473524 geneID 119919435; AMHY Y3 (NC_052095.1):2,513,435-2,535,704 geneID11946726. Gene maps were generated using the Gene Structure Display Server 2.0^60^. To identify transposable elements present in AMHX, AMHY, and the flanking regions of these genes, we used BLASTN^61^ (-task dc-megablast) to search for previously described monotreme TEs from the Repbase database^62^. Sequence alignments and the Phylogenetic trees (Geneious Tree Builder; Genetic distance model: Jukes-Cantor; Tree build method: UPGMA; Resampling method: Bootstrap; 500 replicates) were generated using the Geneious Prime 2022.0.1 and FigTree v1.4.4, dS and dN values were calculated using SNAPv2.1.1 (www.hiv.lanl.gov^63^) Distance was calculated using the DIVEIN (indra.mullins.microbiol.washington.edu/DIVEIN^64^). The non-synonymous (dN) to synonymous (dS) rate ratio (dN/dS) between platypus AMHX and AMHY, or between AMHX or AMHY and AMH(s) in other 10 species, including Alligator, Chicken, Echidna, Human, Koala, Mouse, Opossum, Pig, Wallaby and Zebrafish, were calculated using two different tools, including KaKS_calculator (v2.0)^65^ and PAML (v4.10.6)^66^, independently. Only coding sequences of AMHs were included for the dN/dS calculation. “Model Averaged” method was used for KaKS_calculator and “Nei-Gojobori method” was used for PAML^67^. Proteins were modelled at the Phyre2 server (sbg.bio.ic.uk/∼phyre2^38^) Protein structures were predicted using the Phyre2 server^38^ and AlphaFold multimer^39^ and images produced with ChimeraX (cgl.ucsf.edu/chimerax). Promoter alignments were produced with Clustal Omega (ebi.ac.uk/Tools/msa/clustalo/ ^68^ and transcription factor binding sites identified with MatInspector (genomatix.de^69^) and CIS-BP Database (cisbp.ccbr.utoronto.ca).

### Animals

Adult male and female short-beaked echidna (*Tachyglossus aculeatus*) testis and ovaries were collected opportunistically from injured animals brought into the Currumbin Wildlife Hospital, Queensland, Australia that required euthanasia for animal welfare reasons. Fetal and pouch young echidna samples were collected from a research breeding colony at Currumbin Wildlife Sanctuary (CWS), Queensland, Australia. Fetal samples were dated based on the date of egg laying (oviposition, designated d0 E), with samples varying in age from day 6 to day 9 after oviposition (hatching occurs on day 10) (d6 n=1, d7 n=2, d8 n=3, d9 n=1). Pouch young (py) samples were dated based on the day of hatching (where day of hatching is designated d0 py) (d0 n=3, d2 n=2, d3 n=1, d4 n=1, d9 n=3, d11 n=2). Tissues were collected from wild-caught adult male platypuses and echidnas brought to the Currumbin Wildlife Hospital, Queensland, Australia that required euthanasia for animal welfare reasons. Tissues collected were either stored in RNAlater or fixed in 4% paraformaldehyde and washed ingraded changes of saline:methanol before storing in methanol at 5°C. Wild short-beaked echidnas were obtained and maintained under the Queensland Government EPA scientific purposes permit (WISP153546614). The University of Queensland Animal Experimentation Ethics Committees approved all sampling for echidnas and the University of Melbourne Animal Experimentation Ethics committee approved all sampling of platypuses, in accordance with the National Health and Medical Research Council of Australia Guidelines (2013)^70^.

### Gonad histology

Briefly, tissues were fixed in 4% paraformaldehyde overnight at 4°C, rinsed several times with 1X phosphate buffered saline (PBS), embedded in paraffin and sectioned at 5 μm. After dewaxing and rehydration, slides were stained with Haematoxylin & Eosin according to standard procedures. Sections were examined using an Olympus BX51 microscope and photographed using the attached Olympus DP70 camera (Olympus Corporation).

### Fetal and pouch young gene expression

Total RNA was extracted from pouch young liver, kidney and gonads where available and from adult testis and ovaries with the GenElute Mammalian Total RNA Miniprep Kit (Sigma) and DNase treated using DNA-free (Ambion) according to the manufacturer’s instructions. RNA was then reversed transcribed using the Superscript IV kit (Invitrogen) with oligo(dT) priming according to the manufacturer’s instructions. For all other fetuses and pouch young, gDNA was extracted from formalin fixed tissues with the ReliaPrep FFPE gDNA Miniprep System (Promega) according to manufacturer’s instructions. Adult testis and ovary gDNA was extracted with the Wizard Genomic DNA Purification Kit (Promega) according to manufacturer’s instructions. To identify the sex of formalin-fixed fetuses and pouch young younger than d4 py, RT-PCR was first carried out as previously described^31^ using the male-specific markers *CRSPY* and the RADSeq marker, *RADseq116* on the gDNA samples with *GAPDH* used to confirm the presence of echidna gDNA. To identify the sex of the pouch young where RNA was available RT-PCR was first carried out using the male-specific marker *CRSPY* on the pouch young liver and kidney with adult testis and ovary as positive controls and *GAPDH* as the loading control. RT-PCR was then used to examine the expression profiles of *AMHR2*, *SF-1*, *GATA1*, *GATA4* and *WT1* across the pouch young gonads from d0-d4 py with *GAPDH* as the reference gene and adult testis as the positive control. PCR reactions were amplified with an initial denaturation 95 °C for 5 min, followed by 39 cycles of denaturation at 95 °C for 30 sec, annealing at either 56°C (*CRSPY*) or 60°C (all remaining genes) for 30 sec, and extension at 72 °C for 30 sec, followed by a final extension at 72 °C for 5 min.

### Gene cloning and adult expression

Genes were cloned by RT-PCR from platypus or echidna testis cDNA *AMHY* and platypus ovary cDNA *AMH*. RNA was prepared using TRIzol Reagent (Ambion) and reverse transcription carried out with Superscript III First-Strand Synthesis System (Invitrogen) (for cloning) or iScript cDNA Synthesis Kit (Bio-Rad) (for expression analysis). PCR was carried out with One Taq DNA polymerase (NEB, cloning) using primers listed in Supplementary Table 5 and products cloned using the pGEM-T Easy Vector System I (Promega) or Phusion HF DNA polymerase (NEB) expression analysis.

### *In situ* hybridisation

Regions of low sequence identity between the *AMH* and *AMHY* cDNAs of approximately 500 bp in length were identified, amplified by PCR and subcloned using the pGEM-T Easy Vector System I for use in *in situ* hybridization analysis. The cDNA of platypus *SOX9* in pUCIDT-Amp was obtained from IDT and subcloned into pGEM-T Easy following removal with *EcoRI* (PCR product 499bp) for use in *in situ* hybridization analysis. Clones were linearised and *in vitro* transcribed using SP6/T7 Transcription Kit with Digoxigenin-11-UTP, and detection carried out using NBT/BCIP stock solution (Roche). The *SOX9* clone was linearised with SacII or PstI and transcribed with SP6 or T7 RNA polymerase respectively to generate sense and anti-sense *in situ* probes. The *AMHX* and *AMHY* clones were linearised with NcoI or PstI and transcribed with SP6 or T7 RNA polymerase respectively to generate sense or anti-sense *in situ* probes. *In situ* hybridizations were carried out using a standard protocol. Briefly, tissues were fixed in 4% paraformaldehyde overnight at 4°C, rinsed several times with 1X phosphate buffered saline (PBS), embedded in paraffin and sectioned onto polylysine slides (Menzel-Glaser, Braunschweig, Germany). After dewaxing and rehydration, the sections were washed several times with 1X PBS, glycine, Triton X-100, and triethanolamine buffer and were then immediately hybridized with probes for 16-18 hours at 42°C. Hybridization signals were detected using anti-Dig alkaline phosphatase-conjugated antibody and visualized with NBT/BCIP chromogen, according to the manufacturer’s instructions (Roche GmbH). The sections were counterstained with 0.1% Fast Red (Aldrich Chemical Corp., Milwaukee, WI, USA).

### Chicken Methods

To use the developing chicken embryo as an AMH responsive sex-differentiation model system, the integrating retroviral vector RCASBP was used to overexpress platypus AMH. RCASBP(A) constructs were prepared that express GFP and used a T2A-linker to also express AMH (such that GFP acts as a marker of AMH expression). The “empty” negative control construct encodes just a downstream FLAG-tag (GFP-T2A-FLAG). The chicken AMH ORF was used as a positive control, with known effects^47^ on the chicken urogenital system (GFP-T2A-cAMH). The platypus AMHX and AMHY ORFs were cloned with (+CC) and without the chicken specific cleavage sites (GFP-T2A-pAMHX, GFP-T2A-pAMHX+CC, GFP-T2A-pAMHY, GFP-T2A-pAMHY+CC). Constructs were assembled using homology based cloning methods (HiFi DNA Assembly, NEB Australia) with a combination of synthesized DNA and PCR products with primers to introduce overhangs or sequence modifications. Sanger sequencing was used to confirm correct construct assemblies (Micromon Genomics, Monash University, Clayton Australia).

High titre (> 1 x 10^7^ infectious units per mL) purified virus preparations were made by transfecting each RCASBP construct into DF-1 cells (chicken fibroblast cell line), collecting conditioned media and concentrating using ultracentrifugation as previously described^50^. Purified virus particles were injected into the blastoderms of embryonic day 0 (E0) fertile SPF chicken eggs (Australian SPF Services) and incubated in a humidified environment at 37°C with hourly rocking until E9.5. Eggs were then opened and embryos dissected to expose the urogenital systems for imaging on a Zeiss SteREO Discovery.V12 microscope with Axiocam506Colour camera.

---

**Supplementary Table 1.** Percentage identities of *AMH* coding sequences (base pairs)

|  | Human | Mouse | Opossum | Koala | Wallaby | Echidna_X | Platypus_X | Echidna_Y | Platypus_Y | Chicken |
| --- | --- | --- | --- | --- | --- | --- | --- | --- | --- | --- |
| Human |  | 75 | 61 | 61 | 61 | 60 | 61 | 52 | 52 | 53 |
| Mouse | 75 |  | 57 | 59 | 58 | 58 | 58 | 52 | 52 | 50 |
| Opossum | 61 | 57 |  | 82 | 82 | 65 | 64 | 56 | 57 | 55 |
| Koala | 61 | 59 | 82 |  | 88 | 65 | 65 | 58 | 59 | 55 |
| Wallaby | 61 | 58 | 82 | 88 |  | 65 | 65 | 58 | 59 | 55 |
| Echidna_X | 60 | 58 | 65 | 65 | 65 |  | 93 | 62 | 62 | 57 |
| Platypus_X | 61 | 58 | 64 | 65 | 65 | 93 |  | 61 | 61 | 57 |
| Echidna_Y | 52 | 52 | 56 | 58 | 58 | 62 | 61 |  | 90 | 50 |
| Platypus_Y | 52 | 52 | 57 | 59 | 59 | 62 | 62 | 90 |  | 50 |
| Chicken | 53 | 50 | 55 | 55 | 55 | 57 | 57 | 50 | 50 |  |

**Supplementary Table 2.** Amino acid substitution rates of monotreme AMH proteins.

|  | dS | dS SD | dN | dN SD | Distance |
| --- | --- | --- | --- | --- | --- |
| Platypus_X/Y | 1.66 | 0.24 | 0.35 | 0.12 | 0.56 |
| Echidna_X/Y | 1.55 | 0.21 | 0.32 | 0.10 | 0.58 |
| Platypus_X/Echidna_X | 0.18 | 0.02 | 0.04 | 0.01 | 0.08 |
| Platypus_Y/Echidna_Y | 0.25 | 0.03 | 0.07 | 0.01 | 0.11 |

**Supplementary Table 3.** Percentage identities of AMH proteins.

| Species | Platypus_X | Platypus_Y | Echidna_X | Echidna_Y |
| --- | --- | --- | --- | --- |
| Alligator | 36 | 34 | 36 | 36 |
| Chicken | 36 | 34 | 36 | 35 |
| Lizard | 38 | 34 | 37 | 35 |
| Koala | 50 | 48 | 50 | 48 |
| Wallaby | 50 | 48 | 50 | 48 |
| Opossum | 50 | 46 | 50 | 47 |
| Human | 43 | 40 | 42 | 41 |
| Pig | 42 | 39 | 42 | 40 |
| Mouse | 39 | 38 | 38 | 39 |
| Pejerrey | 15 | 17 | 16 | 17 |
| Pejerrey_Y | 16 | 17 | 16 | 18 |
| Tilapia | 15 | 16 | 15 | 16 |
| Tilapia_Y | 15 | 16 | 15 | 16 |
| Zebrafish | 17 | 18 | 16 | 17 |
| Platypus_X | 100 | 52 | 91 | 52 |
| Echidna_X | 91 | 52 | 100 | 52 |
| Platypus_Y | 52 | 100 | 52 | 86 |
| Echidna_Y | 52 | 86 | 52 | 100 |

**Supplementary Table 4.**
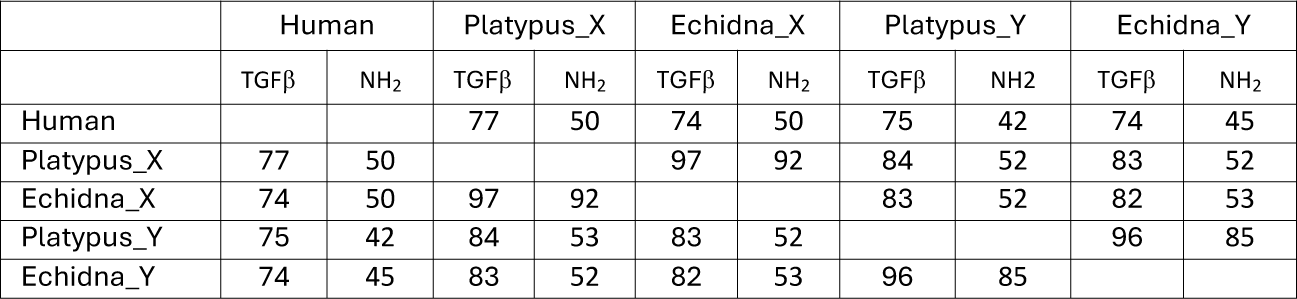
Percentage identities of TGF-β and NH2 domains of AMH proteins.

**Supplementary Table 5.**
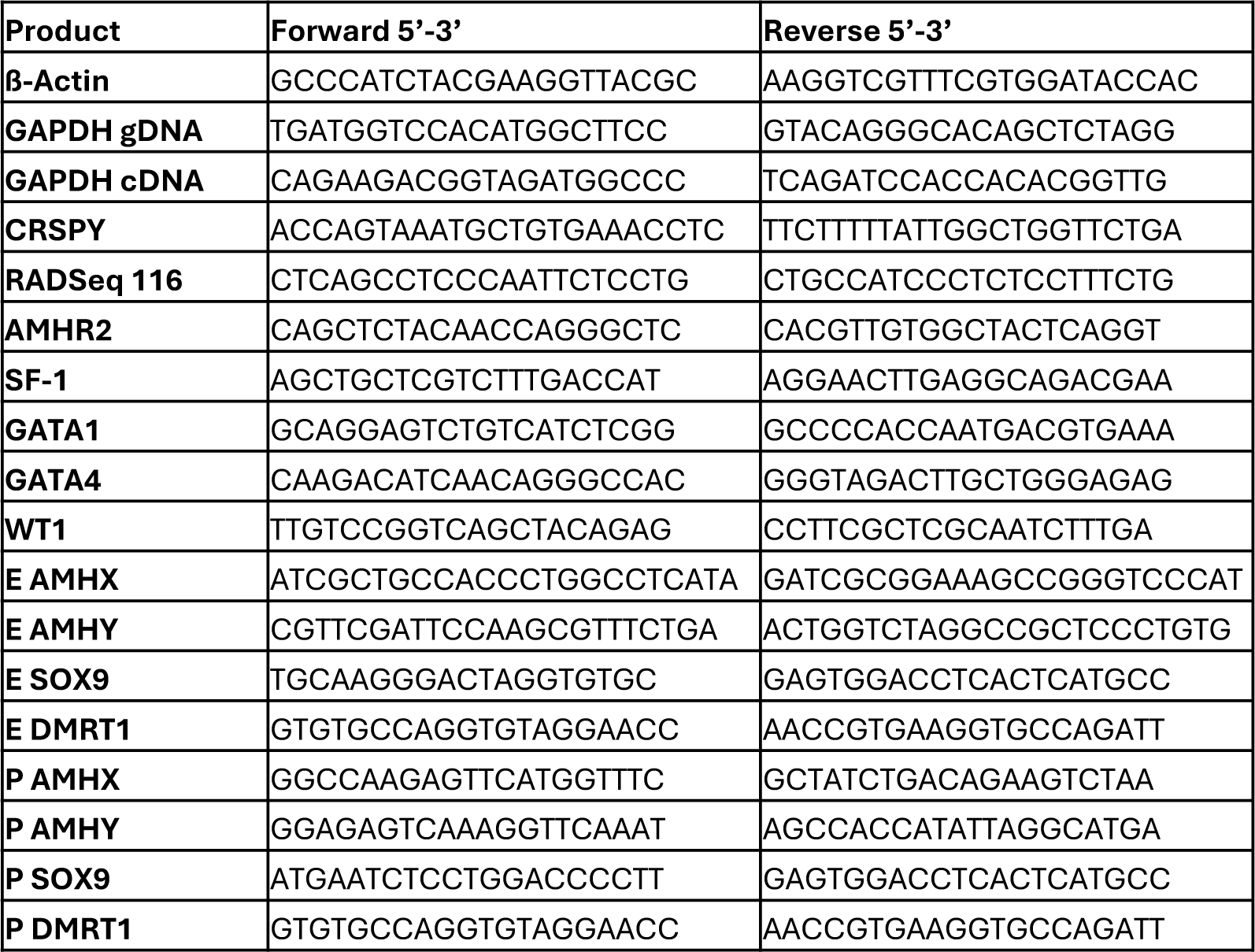
Primers used for RT-PCR and generation of *in situ* hybridization probes.

**Supplemental Data Fig. 1.**
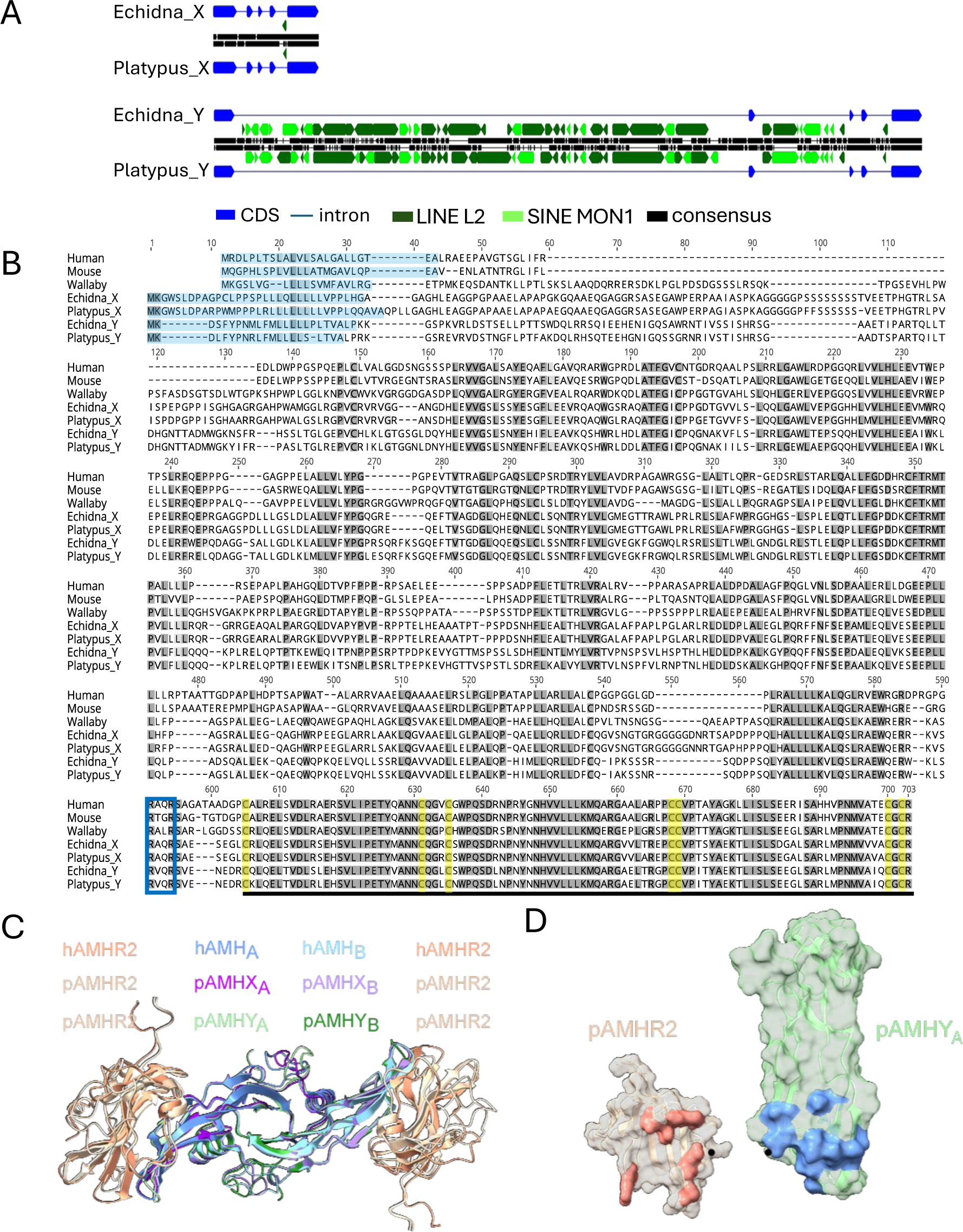
Analysis of monotreme AMH genes and proteins. (A) Repeat content of the monotreme *AMHX* and *AMHY* genes. Coding sequence, blue; introns, black or blue line, LINE L2, dark green and SINE MON1, light green. (B) Alignment of AMH proteins from mammalian species human and mouse (eutherian), wallaby (metatherian), echidna and platypus (prototherian). Shaded amino acids, identical amino acids; light blue shading, predicted signal peptide; yellow shading; conserved cysteine residues; blue outline, cleavage site between amino terminal and TGF-β domains and underline, TGF-β domain. (C) Overlay of platypus AMHR2-AMHX and AMHR2-AMHY multimeric complexes, predicted with AlphaFold multimer (Evans et al., 2021), using a stoichiometry of 2 AMHR2 domains and 2 AMHX (or Y) domains. The experimentally determined human hAMHR2-AMH structure (PDB 7l0J) is also shown in the overlay. (D) Pull apart view of the predicted platypus AMHR2 (peach) and AMHY (pale green) interface to highlight close contact residues conserved between the platypus and human complexes. Close contact residues from the interface in the hAMHR2-AMH complex (PDB 7l0J) are shown highlighted in the platypus cAMHR2-AMHY complex if the residue is identical (salmon in AMHR2; pale blue in AMHY).

**Supplemental Data Fig. 2.**
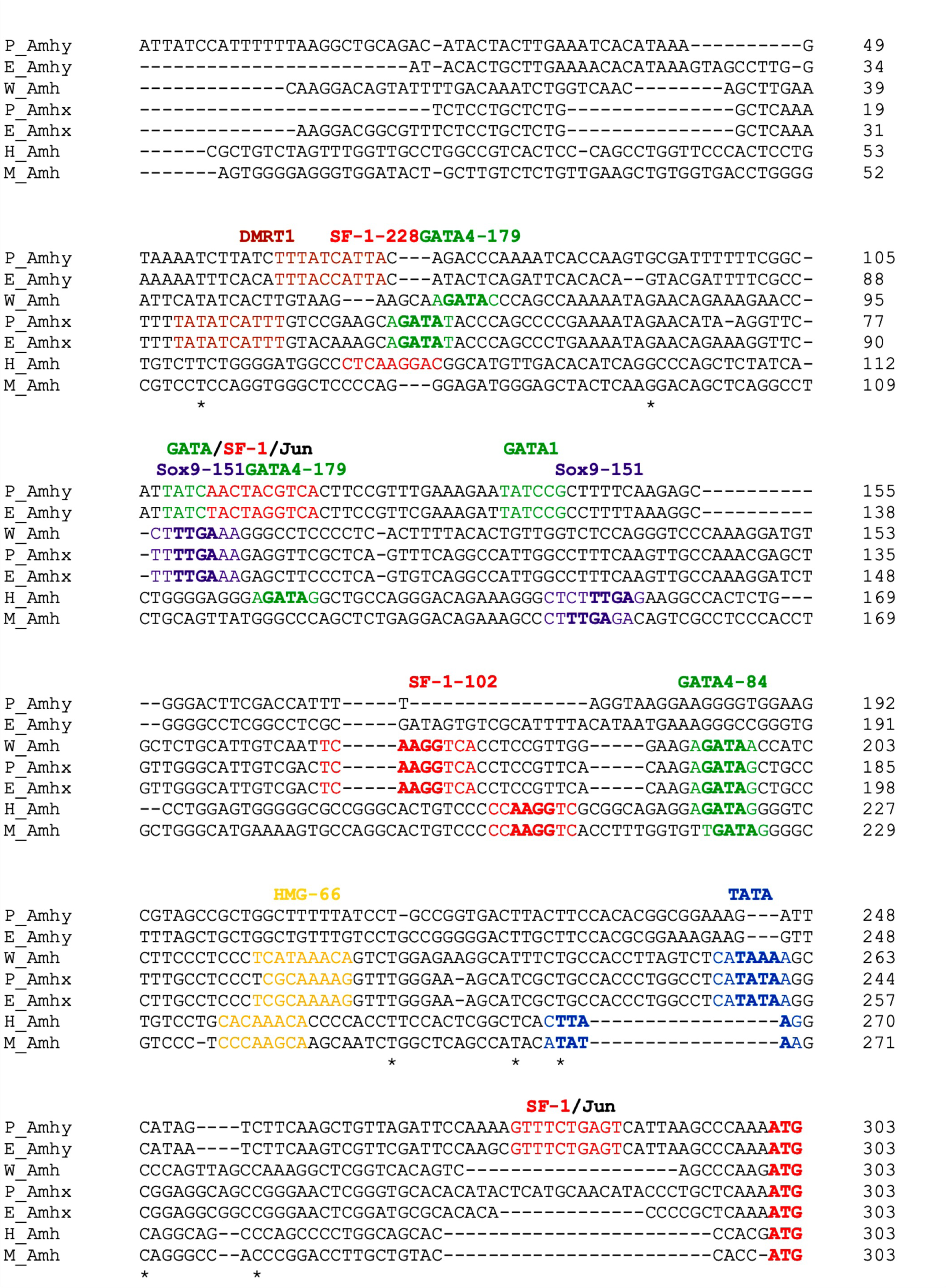
Mammalian *AMH* proximal promoter alignment 300 bp upstream of the translation start site, showing putative transcription factor binding sites and relative positions on the human promoter. P, platypus; E, echidna; W, wallaby; H, human and M, mouse; DMRT1 binding site, brown; GATA4 and GATA1 binding site, green; SF-1 and SF-1/JUN binding site, red; SOX9 binding site, purple; HMG binding site, yellow and TATA, binding site, blue.

**Supplemental Data Fig. 3.**
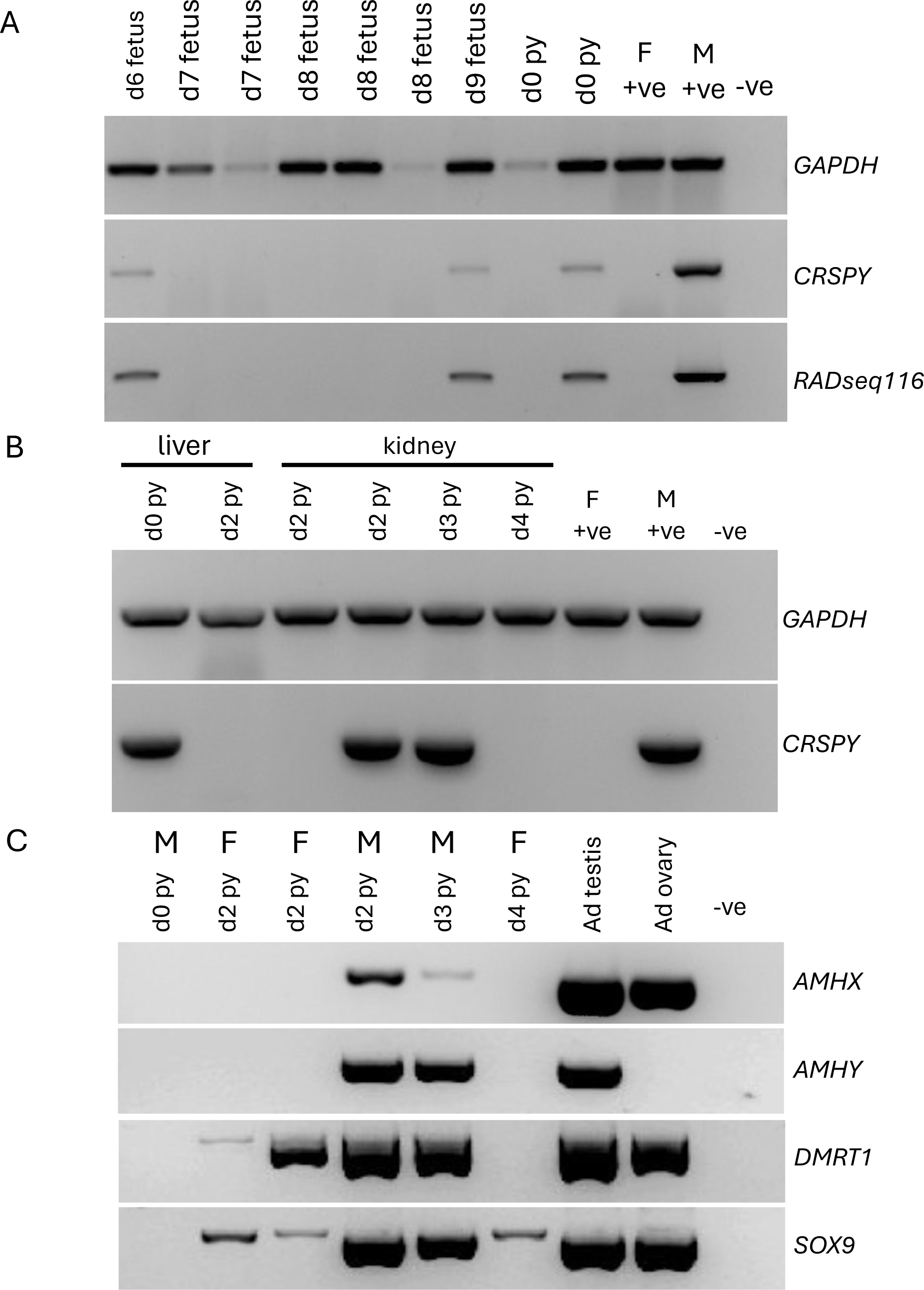
Echidna sexing and expression analysis by RT-PCR. Expression of *CRSPY* and *RADSeq 116* with *GAPDH* as loading control detected by RT-PCR from either FFPE gDNA (A) or somatic pouch young tissues (B) to identify the sex of all fetuses and pouch young with adult testis and ovary used as male and female positive controls. (C) Expression of *AMHX*, *AMHY, DMRT1* and *SOX9* in pouch young gonads from d0-d4 py with adult testis and adult ovary as a positive control. Py, pouch young; Ad, adult.

**Supplemental Data Fig. 4.**
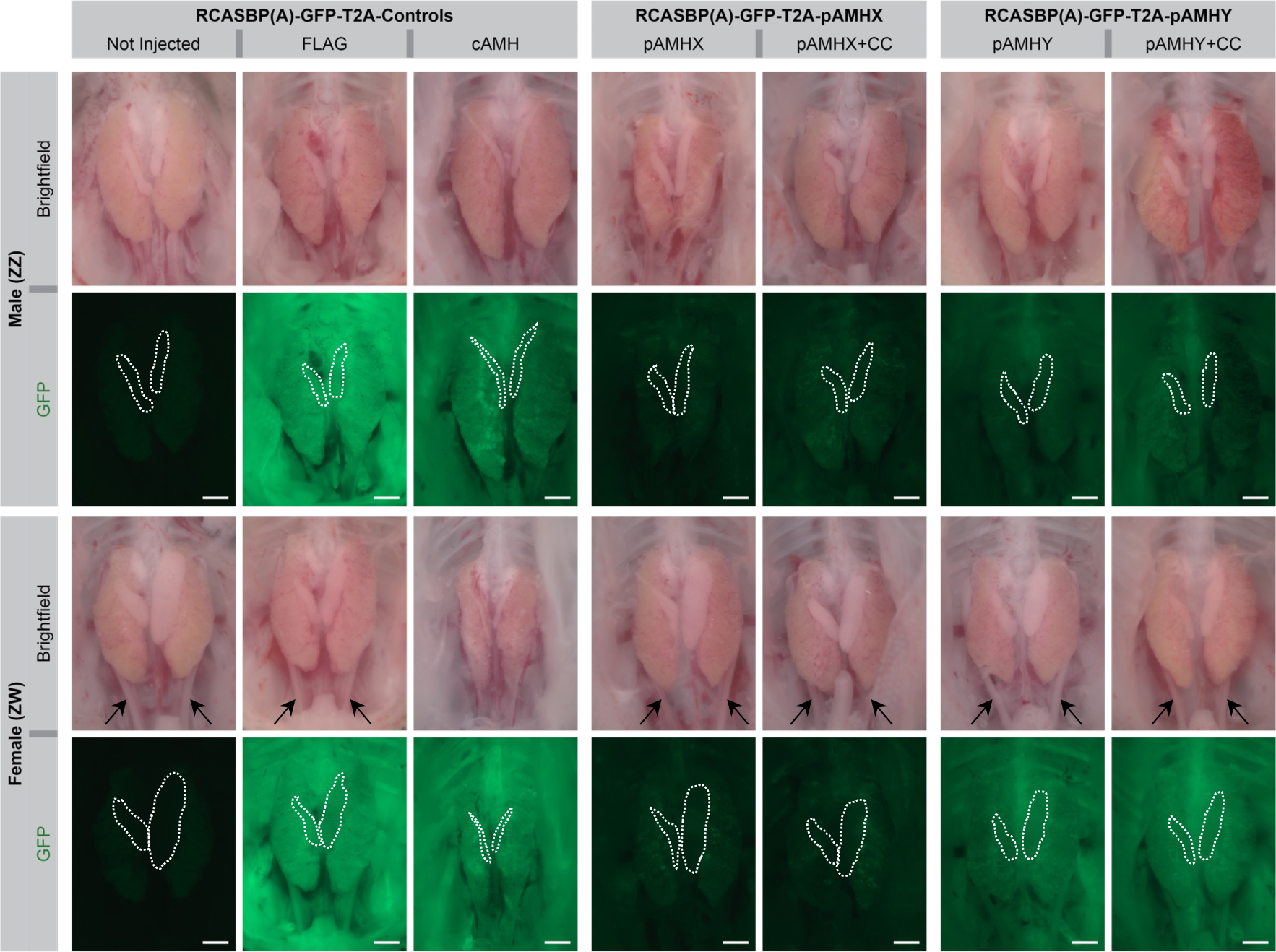
Overexpression of platypus AMHX and AMHY in the chicken. Dissected urogenital systems at E9.5 from genetically male and female chicken embryos (determined by PCR). GFP signal indicates expression of the indicated transgenes (GFP-T2A-ORF, platypus proteins, +CC carry the chicken cleavage site). The gonads are outlined (white dotted lines) with the Müllerian ducts where present indicated (black arrows). Scale bar, 1mm.

**Supplemental Data Fig. 5.**
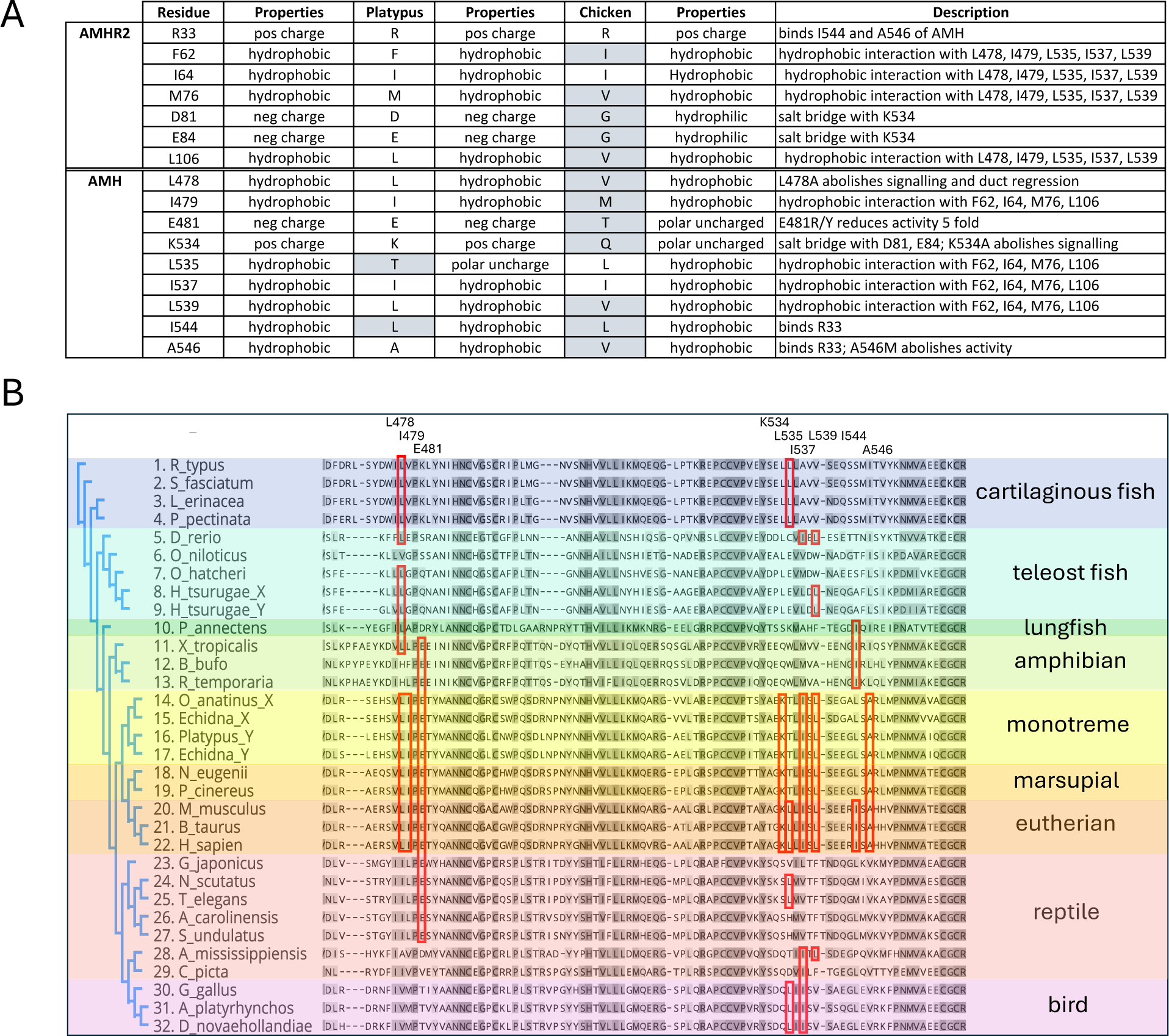
Comparative analysis of residues involved in AMH binding to AMHR2. (A) Table showing a comparison of AMHR2 and AMH amino acids involved in receptor binding between human, platypus and chicken. Residues not conserved with human, grey shading. (B) Alignment of C-terminal of the AMH TGF-β domain protein sequences from vertebrates showing conservation of amino acids involved in receptor binding defined in the human protein (Hart *et al*., 2021, red boxes).

## Acknowledgments

Particular thanks to the veterinary staff and keepers at Currumbin Wildlife Sanctuary for assistance with echidna monitoring. This project was supported by an Australian Research Council Discovery Project grant to FG and Linkage grant to MR, SJ and MP.

## Author contributions

LS-W and FG designed the experiments to characterise the monotreme *AMH* genes and proteins. LS-W cloned and sequenced the monotreme *AMH* genes, carried out synteny analysis, gene analysis, protein promoter analysis and prepared the *in situ* hybridisations of platypus samples. JF, SJ, MP and MR designed the echidna experiments. MR and JF collected the fetal and pouch young echidna samples and MP supervised the echidna colony. MR, JF and SJ dissected the gonad samples. JF prepared the fetal and pouch young gonad RT-PCRs. HY and JF prepared the *in situ* hybridisations of echidna samples. YZ and GZ provided the monotreme genomic sequences of *AMHX* and *AMHY*. LS-W, FG, AM and CS designed the experiments in the chicken model system, with AM and CS performing the *in ovo* work and data acquisition. ZQ carried out the Ka/Ks analysis. KS carried out the AMH and AMHR2 protein modelling and analysis. JG, AS and DA carried out the repeat analysis. LS-W, JF, CS, MR and FG wrote the manuscript. All authors revised the manuscript and approved the paper.

